# Amyloid precursor protein: an Achilles’ heel to exploit in the treatment of prostate cancer

**DOI:** 10.64898/2026.09.21.752657

**Authors:** Mai Alhadrami, Emily Gill, Claire M Perks, Rachel M Barker

**Author notes:** **Corresponding Author:** Claire M Perks, Professor of Cancer Endocrinology, Bristol Medical School, Translational Health Sciences, Learning & Research Building, Cancer Endocrinology Group, Southmead Hospital, Bristol, BS105NB. co-senior authors (Claire M Perks, Rachel M Barker).

## Abstract

**Background:** Epidemiological studies have highlighted an inverse association between cancer and Alzheimer’s disease (AD), but the mechanisms underlying this association are not currently understood. Amyloid precursor protein (APP) and its cleavage may represent one potential mechanism. Its importance has been heavily researched due to it giving rise to the production of amyloid-β, a pathological hallmark of AD. APP, however, has not been extensively investigated in cancer.

**Methods:** FFPE prostate tumour and adjacent benign tissue immunohistochemically stained for APP, androgen receptor and the insulin-like growth factor-I receptor (IGF1R). Normal prostate PNT2, androgen sensitive LNCaP and androgen insensitive PC3 *in vitro* were used to assess the effects of androgen, IGF-I and their inhibitors on APP and its cleavage; changes in protein abundance were measured by western blot; interactions between proteins using co-immunoprecipitation and effects on cell proliferation, apoptosis and cell signalling of *APP* silenced cells using a BrdU proliferation assay, Muse Cell Analyser and by TMT proteomic analysis respectively.

**Results:** Data showed that APP was upregulated in prostate tumour tissue compared to adjacent benign. Furthermore, in cancer cell models, in contrast to normal prostate epithelial cells, it was observed that APP was critical for prostate cancer cell survival by interacting with androgen and IGF-I signalling pathways.

**Conclusions:** Drugs that target APP have already been developed for AD and our data suggests they offer an exciting opportunity for repurposing in the treatment of prostate cancer.

## Introduction

Prostate cancer is the second most commonly diagnosed cancer worldwide and the fifth leading cause of cancer-related death, with an estimated 397000 prostate cancer deaths in 2022 [1]. Despite major advances in prostate cancer treatment over the last few decades, important challenges remain, including prevention of progression to castrate-resistant prostate cancer (CRPC) and overcoming development of therapy resistance [2, 3], highlighting the need for new therapeutic strategies.

Alzheimer’s disease (AD) is a progressive neurological condition that irreversibly impairs memory and cognition, affecting over 47 million people worldwide [4]. Aging is the primary risk factor for both prostate cancer and AD, and the two diseases also share other risk factors, including obesity and Type 2 Diabetes, however, interestingly, numerous population studies have shown an inverse association between the two [5–7]. The mechanisms underlying this inverse relationship remain poorly understood, however amyloid precursor protein (APP) and its cleavage products may be one potential candidate underpinning this inverse relationship. APP and its processing has been extensively investigated in AD as its cleavage by β-secretase, BACE1, followed by γ-secretase gives rise to the production of amyloid-β (Aβ) peptides, that aggregate to form amyloid plaques, a key neuropathological hallmark of AD [8]. This amyloidogenic cleavage pathway also gives rise to the production of soluble APPβ (sAPPβ). APP may alternatively be cleaved by α-secretases, including A disintegrin and metalloproteinase (ADAM)-10 and -17 (also known as TNF-α Converting Enzyme, TACE) and BACE2, precluding Aβ production but producing soluble APPα (sAPPα) which has been shown to promote neurogenesis and neuronal survival [8, 9].

Despite APP being abundant in non-neuronal tissues and overexpressed in many cancer types, including prostate [10, 11], its role in cancer has not been extensively studied. In breast cancer, APP, particularly *via* its non-amyloidogenic processing, has been shown to promote cell growth and migration *via* MAPK signalling [12–15]. Similarly, in prostate cancer, APP has been associated with enhanced cell growth and migration [16] and shown to be regulated by androgen [11].

Androgen is an important driver of growth in androgen-sensitive prostate cancer, hence its association with APP may be of importance. Insulin-like growth factor (IGF) signalling is also important in the development and progression of prostate cancer and IGF-I has been associated with more aggressive disease [17]. Whilst IGF-I is neuroprotective and promotes non-amyloidogenic APP cleavage in models of AD [18], the relationship between IGF signalling and APP in cancer remains unclear.

This seminal study encompasses a detailed examination of the role of APP in prostate cancer, assessing its expression in prostate tumour tissue, how androgen and IGF-I, key drivers of prostate cancer, affect APP and its processing, and investigating the effects of silencing the *APP* gene, inhibiting α-secretase cleavage of APP and the effects of addition of exogenous sAPPα.

## Methods

### TCGA analysis

The Cancer Genome Atlas Prostate Adenocarcinoma (TCGA-PRAD) dataset was used to compare expression of the *APP* gene in prostate tumour vs normal tissue and between different Gleason grades in the tumour cohort. Correlations between *APP* and the genes encoding the androgen receptor (AR) and the insulin-like growth factor receptor (IGF1R) were also assessed in the tumour cohort.

### Tissue acquisition and immunohistochemistry

Tissue from the Prostate Cancer: Evidence of Exercise and Nutrition Trial (PrEvENT) (14/SW/0056) was used to assess the abundance of APP in tumour and surrounding benign tissue. Details of this cohort have been published previously [19, 20]. Tissue was formalin fixed and paraffin embedded (FFPE), cut to 4μm and collected on adhesive slides. Immunohistochemical staining was performed using a Ventana BenchMark Ultra Immunostainer system (Ventana Medical Systems) according to the manufacturer’s protocol. Briefly, tissue sections were deparaffinised then subjected to citrate antigen retrieval before incubation with an APP antibody (1:4000, Abcam) for 2 h. Slides were counterstained with haematoxylin for 8 min and with bluing reagent for 4 min (Ventana Medical Systems), dehydrated and mounted in Clearium mounting media (Leica Biosystems). An Allred scoring system was used to assess APP staining as described previously [21].

### Cell Culture

The human prostate cancer cell lines LNCaP and PC3 were purchased from the American Type Culture Collection (ATCC, Middlesex, UK). The normal epithelial prostate cell line, PNT2, was purchased from ECACC (London, UK). Authentication was confirmed using short tandem repeat (STR) analysis, and routine testing ensured they were mycoplasma-free. Cells were cultured as described previously [22]. Briefly, PNT2 and LNCaP cells were maintained in RPMI1640 medium and PC3 cells were maintained in DMEM, all supplemented with 10% foetal bovine serum (FBS) and 1% L-glutamine. Cells were passaged upon reaching 80% confluence. For experiments with dihydrotestosterone (DHT) or IGF-I treatment, cells were serum starved for 24 h and dosed in serum free media; all other treatments were carried out in growth media. Cell treatments included DHT (Sigma, 0.001-0.1μM), anti-androgen, enzalutamide (Sigma, 1-5μM), IGF-I (Gropep, 25-100ng/mL), an IGF-IR inhibitor, AG1204 (1-3uM), sAPPα (2-8nM) and an ADAM-10/ADAM-17 dual inhibitor, GW 280264X (Tocris Bioscience, 10-1000nM).

### RNA silencing

*APP* was silenced using siRNA (SI02780288, Qiagen) and cells were transfected using Lipofectamine® RNAiMAX (Invitrogen). AllStars Negative Control siRNA (Qiagen) was used as the control. Cells were seeded in 6 well plates at a density 2-5×10^5^ cells per well and incubated with 30nM or 60nM siRNA in Opti-MEM (Thermofisher) for 48 hr. Successful *APP* silencing was confirmed by western blot.

### Tritiated thymidine incorporation assay (TTI)

Tritiated thymidine incorporation assay was used to assess proliferation, as described previously [23]. Briefly, cells were seeded in 24 well plates at a density of 2 x10^5^ cells per well, then incubated with 0.1µCi [³H]-thymidine for the final 4 hours of the treatment period. Following incubation with 5% trichloroacetic acid (Merck) then sodium hydroxide (Fisher Scientific), the cell suspension was then transferred to tubes containing scintillation fluid and disintegrations per minute (DPM) were read on a Beckman Scintillation Counter LS6500.

### BrdU (5-bromo-2’-deoxyuridine) incorporation assay

A BrdU incorporation assay kit (Roche) was used to assess proliferation, as per the manufacturer’s protocol. Briefly, cells were seeded in 96 well plates at a density of 5-10×10^3^ cells per well and incubated with BrdU label for the last 2 hours of the treatment period. Cells were fixed and then incubated with the anti-BrdU-peroxidase antibody, followed by incubation with TMB (3,3’,5,5’-tetramethylbenzidine) substrate. Absorbance was measured at 450nm using a plate reader (BioRad).

### Cell viability and apoptosis assessment using the Muse Cell Analyser

For cell counting and apoptosis experiments, cells were seeded in 6 well plates at a density of 2-5×10^5^ cells per well. For cell counting, the Muse Count and Viability kit (MCH100102, Cytek Biosciences) was used, as per the manufacturer’s instructions. Following treatments, cells were collected and resuspended in growth media at a concentration of 1×10⁵ to 1×10^7^ cells/ml, then diluted 1:10 with the Count and Viability Reagent for 5 min before loading into the Muse Cell Analyser. Output measures included concentrations (cells/mL) for total, live and dead cell counts. To assess apoptosis, the Muse Annexin V and Dead Cell kit (MCH100105, Cytek Biosciences) was used, as per the manufacturer’s instructions. Following treatments, cells were collected and resuspended in growth media at a concentration of 1×10⁵ to 1×10^7^ cells/ml, diluted 1:1 with Annexin V and Dead Cell reagent and incubated at room temperature in the dark for 20 min before loading into the Muse Cell Analyser. Output measures included concentrations (cells/mL) for live, early apoptotic, late apoptotic and dead cells.

### Western blotting

Western blotting was performed as described previously [24]. Briefly, 30 μg total protein were run on 4-20% SDS-PAGE then transferred to nitrocellulose membrane (BioRad) and immunoblotted with the following antibodies: APP (1:1000, Abcam), ADAM-10 (1:1000, Antibodies), TACE (1:500, Abcam), BACE1 (1:1000, Abcam), BACE2 (Santa Cruz, 1:500), sAPPα (1:1000, BioLegend), sAPPβ (Merck, 1:1000), cleaved PARP (1:1000, BD Pharmingen), AR (1:1000, Cell Signalling), IGF1R (1:1000, Cell Signalling), AKT (1:1000, Cell Signaling), p-AKT (1:1000, Cell Signaling) and GAPDH (1:2500, Merck), following the manufacturer’s instructions. After incubation with appropriate peroxidase-conjugated secondary antibodies (Sigma), proteins were visualised by Clarity ECL substrate (BioRad) using BioRad Chemidoc XRS + system and analysed using Image lab software (BioRad). Supernatants were first concentrated by centrifugation at 8,000 *xg* at 4°C for 20 minutes in Amicon® Ultra Centrifugal Filters prior to western blotting (Merck).

### ELISA

Levels of sAPPα following treatment with GW 280264X were measured in cell supernatants using an ELISA kit (27734, Stratech), as per the manufacturer’s instructions. Briefly, supernatants were diluted 1:10 with enzyme immunoassay (EIA) buffer (provided) and loaded in duplicate with a recombinant protein standard curve and negative controls onto a 96 well plate. Following overnight incubation at 4⁰C, the plate was washed and then incubated with peroxidase-labelled antibody for 30 min at 4⁰C. Following incubation with TMB substrate, the reaction was stopped and absorbance was measured at 450nm using a plate reader (BioRad).

### Proteomics

Tandem-mass tagging proteomics was performed on *APP* silenced LNCaP and PC3 cells at the University of Bristol Proteomics Facility. 100µg total protein at a concentration of 2mg/mL were prepared for each cell lysate. Effective *APP* silencing was confirmed by western blot.

### Co-immunoprecipitation

500μg of cell lysate were incubated with 2μg APP antibody (Abcam) or 2μg rabbit IgG isotype control (Cell Signalling) for 2 h at 4⁰C with constant agitation. 50μl protein A agarose beads (Insight Biotechnology) were added to the lysate-antibody mix and incubated for 2 h at 4⁰C with constant agitation. Samples were centrifuged at 2000 xg at 4⁰C for 1 min and the supernatant removed. Three washes with lysis buffer were performed. Laemmli buffer was added and the samples boiled for 10 min before proceeding to western blot.

### Statistical analysis

Statistical analysis was performed using GraphPad Prism software. Results are expressed as mean ± standard error of the mean (SEM) from three independent experiments, each performed in triplicate. For group comparisons one-way ANOVA was used, followed by Dunnett’s test for comparison to a control group or Bonferroni’s multiple comparisons test for comparison of all groups. Spearman’s correlation was used to assess correlations. A p-value of less than 0.05 (p < 0.05) was considered statistically significant. Statistics were performed on the raw data for graphs showing fold change control.

For analysis of proteomic data, RStudio Bioconductor packages were used. Differentially expressed proteins (DEPs) analysis was performed using thresholds of Log2 fold change (Log2FC) >1 or < -1 and p < 0.05. DEP was visualized using the Enhanced Volcano package. Kyoto Encyclopedia of Genes and Genomes (KEGG) pathway enrichment analysis was performed using the cluster Profiler package. Gene Ontology (GO) enrichment analysis was carried out using enrich plot. Figures were generated using ggplot2.

## Results

### APP is increased in prostate cancer compared to normal prostate tissue

Using tissue from the PrEvENT study, we performed immunohistochemical assessment of APP in in prostate tumour and adjacent benign tissue. APP abundance was significantly increased in tumour tissue compared to benign (p<0.001) (**Fig 1A**) and there was a trend towards increased APP abundance in mid-grade tumours and reduced abundance at higher Gleason grades (**Fig 1B**). Comparison of *APP* gene expression in the TCGA PRAD prostate cancer and the GTex normal prostate mRNA datasets showed a non-significant increase in *APP* expression in tumour tissue compared to normal (**extended data Fig 1A**) and there was a significant reduction in APP expression in Gleason grades of 8 and higher compared to Gleason grades 6 (p<0.001) and Gleason grade 7 (p<0.0001) (**extended data Fig 1B**).

**Figure 1:**
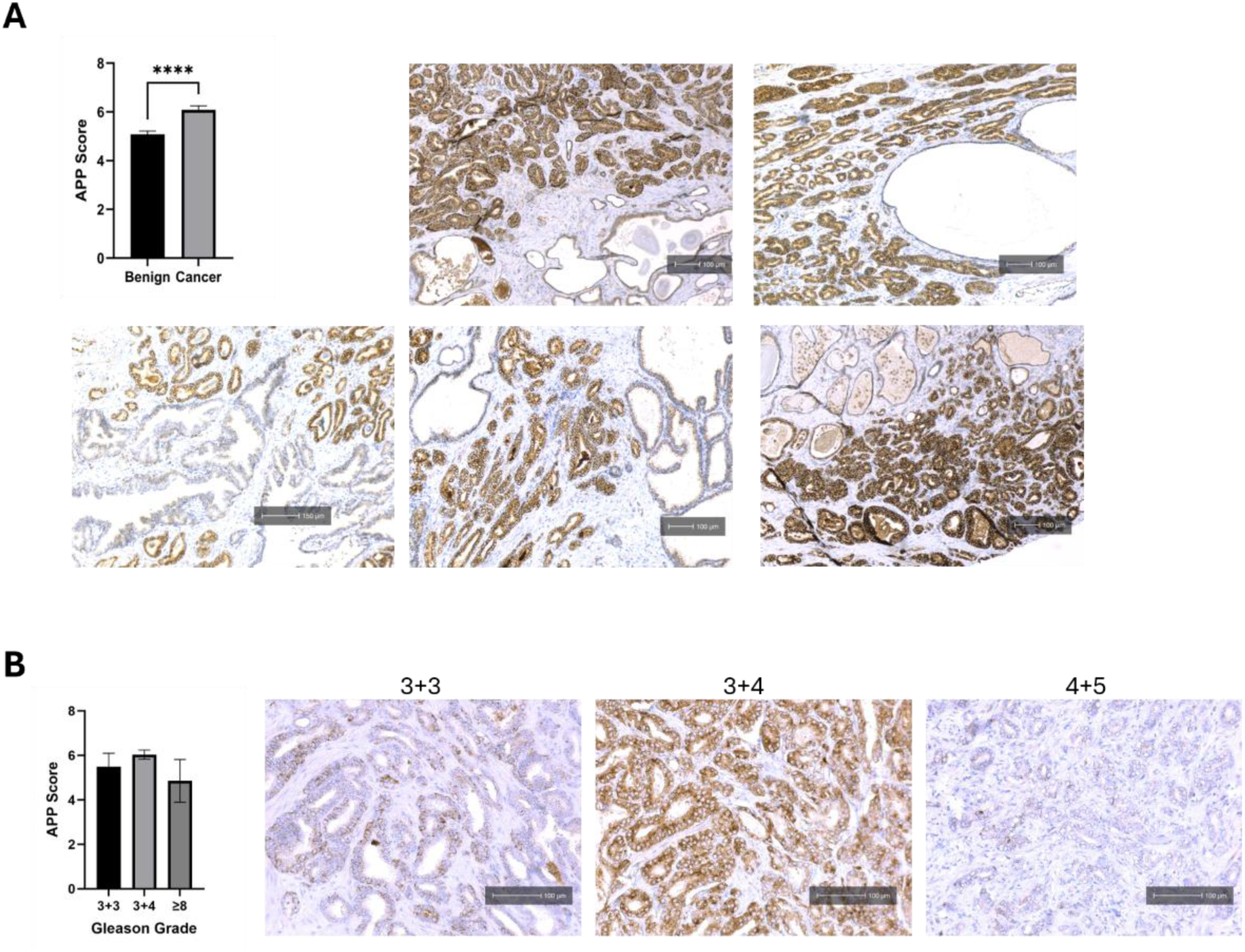
Immunohistochemical analysis of APP in prostate tissue from the PrEvENT study. **(A)** APP abundance was significantly higher in tumour compared to adjacent benign tissue (p<0.0001). Images show representative immunohistochemical staining from 5 patients. Magnification = x10, scale bar = 100-150 μm. **(B)** APP abundance was not significantly changed with respect to Gleason grade, however there was a trend towards reduced APP at higher Gleason grades. Images show representative staining of APP in tumour tissue in a low-, mid –and high-grade sample. Magnification = x20, scale bar = 100μm.

### Drivers of prostate cancer growth affect APP and its processing

#### Effects of androgen on APP and its cleavage in LNCaP cells

In LNCaP cells, DHT treatment caused a dose dependent increase in abundance of APP and α-secretases ADAM10 and TACE that were significant at the 0.1μM DHT dose, which induces the most significant proliferative effect (APP p<0.001, ADAM-10 p<0.001 and TACE p<0.05). BACE2 was also significantly increased at the 0.1μM DHT dose (p<0.001). BACE1 was reduced in response to DHT, but not significantly (**Fig 2A**). The resulting sAPPα fragments produced by cleavage of APP by α-secretases were significantly increased at 0.1μM DHT dose, whilst sAPPβ, produced by BACE1 cleavage, was reduced, but not significantly (**Fig 2B**).

**Figure 2:**
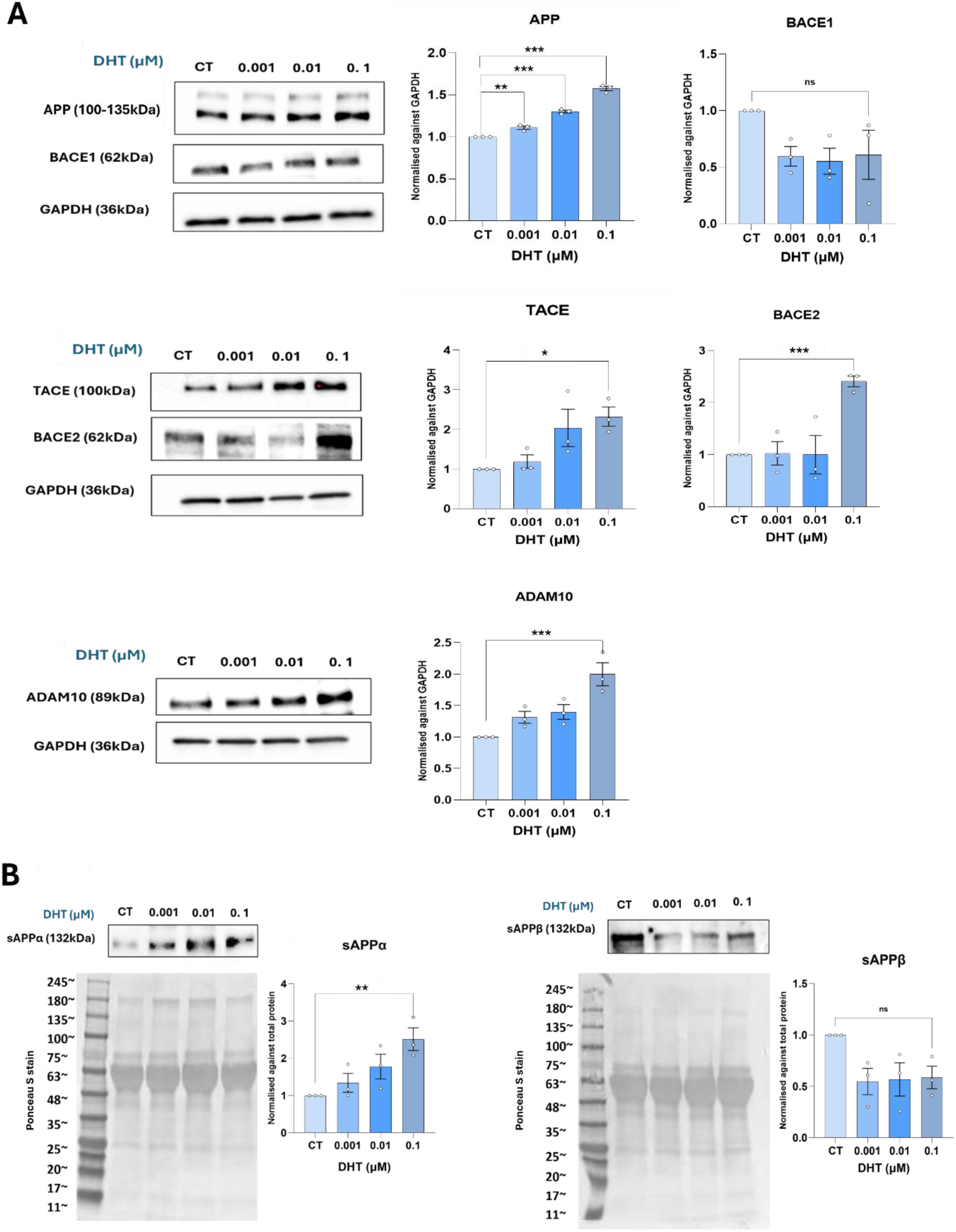
Assessment of the effects of DHT on APP, secretase enzymes and sAPP production in LNCaP cells. **(A)** DHT caused dose-dependent increases in APP, ADAM-10, TACE and BACE2, all of which were significant at the 0.1μM DHT dose (p<0.001, p<0.001, p<0.05 and p<0.001, respectively). In contrast, BACE1 was reduced in response to DHT but not significantly. **(B)** Assessment of sAPPα and sAPPβ secreted into the cell supernatants showed a dose-dependent increase in sAPPα which was significant at the 0.1µM DHT dose (p<0.05). sAPPβ levels were reduced in response to DHT but not significantly. Representative western blots are shown, and graphs represent densitometric analysis of bands corrected to either the GAPDH loading control or Ponceau S stain. Bars represent fold change from control (mean ± SEM, n=3).

#### Effects of anti-androgen enzalutamide on APP and its cleavage

In LNCaP cells, enzalutamide caused dose-dependent reductions in APP (p<0.01) and α-secretases BACE2 (p<0.01) and TACE (p<0.001). The abundance of α-secretase ADAM-10 was not significantly altered by enzalutamide whilst the β-secretase BACE1 was significantly increased at the 5µM enzalutamide dose (p<0.05) (**Fig 3A**). The resulting sAPPα fragments produced by cleavage of APP by α-secretases were significantly reduced (p<0.01), whilst sAPPβ, produced by β-secretase cleavage, was significantly increased (p<0.01) at the 5μM enzalutamide dose (**Fig 3B**).

**Figure 3:**
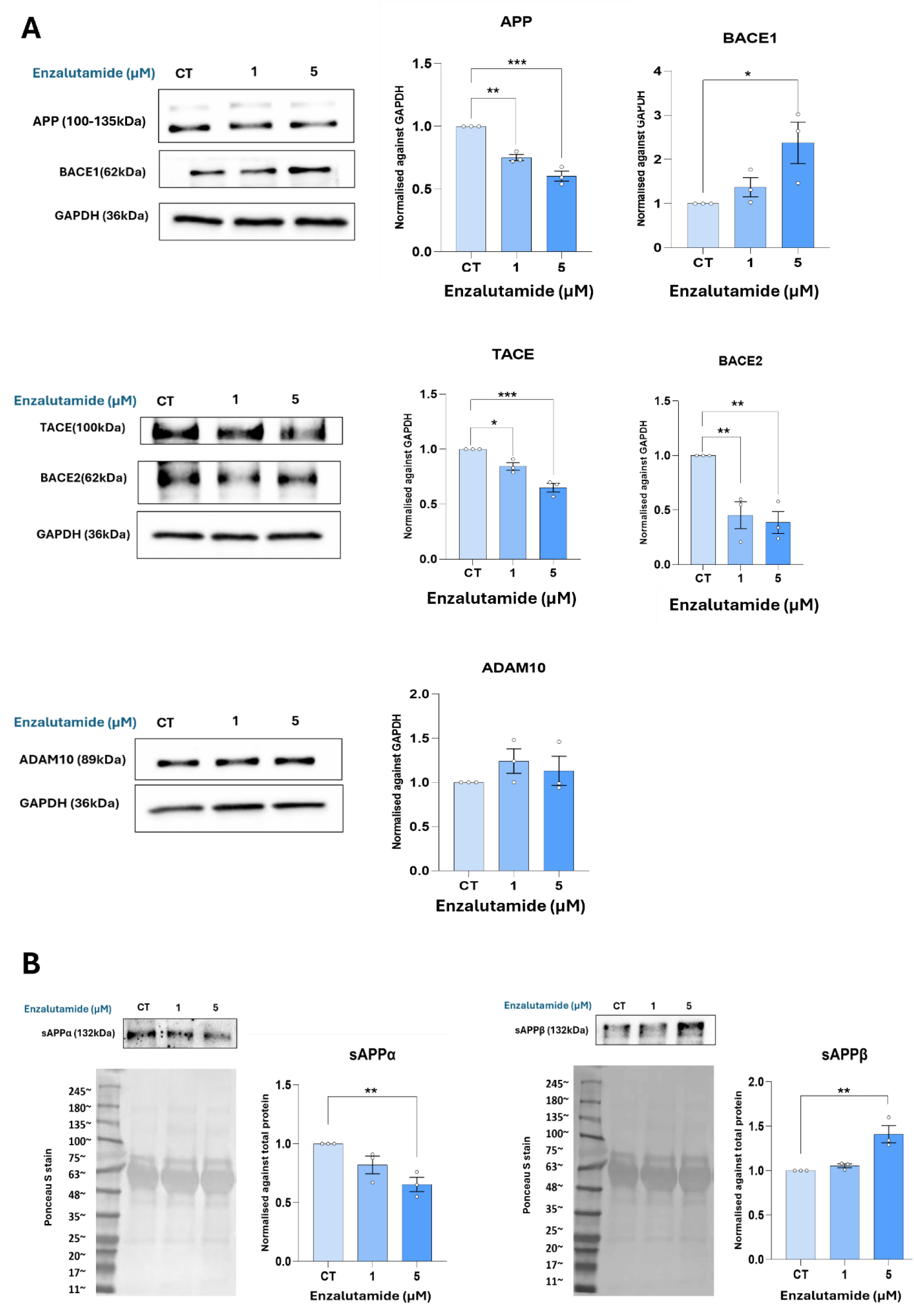
Assessment of the effects of enzalutamide on APP and its secretase enzymes in LNCaP cells. **(A)** Enzalutamide caused dose-dependent reductions in APP (p<0.01), BACE2 (p<0.01) and TACE (p<0.001). ADAM-10 abundance was not significantly changed by enzalutamide whilst BACE was significantly increased at the 5µM enzalutamide dose (p<0.05). **(B)** Assessment of sAPPα and sAPPβ secreted into the cell supernatants showed a dose-dependent reduction in sAPPα and an increase in sAPPβ, both of which were significant at the 5µM enzalutamide dose (p<0.01). Representative western blots are shown, and graphs represent densitometric analysis of bands corrected to either the GAPDH loading control or Ponceau S stain. Bars represent fold change from control (mean ± SEM, n=3).

#### Effects of IGF-I on APP and its cleavage in PC3 cells

In androgen-independent PC3 cells, IGF-I caused a significant increase in abundance of both APP (p<0.05) and the α-secretase BACE2 (p<0.05). ADAM-10 and TACE were not significantly altered by IGF-I, whilst BACE1 was significantly reduced at the 50ng/ml IGF-I dose (p<0.05) (**Fig 4A**). The resulting soluble APP fragments, sAPPα and sAPPβ, were both dose-dependently increased in the cell supernatants, with the increases in sAPPα being significant at all three IGF-I doses (25ng/ml p<0.05, 50ng/ml p<0.001, 100ng/ml p<0.001) (**Fig 4B**).

**Figure 4:**
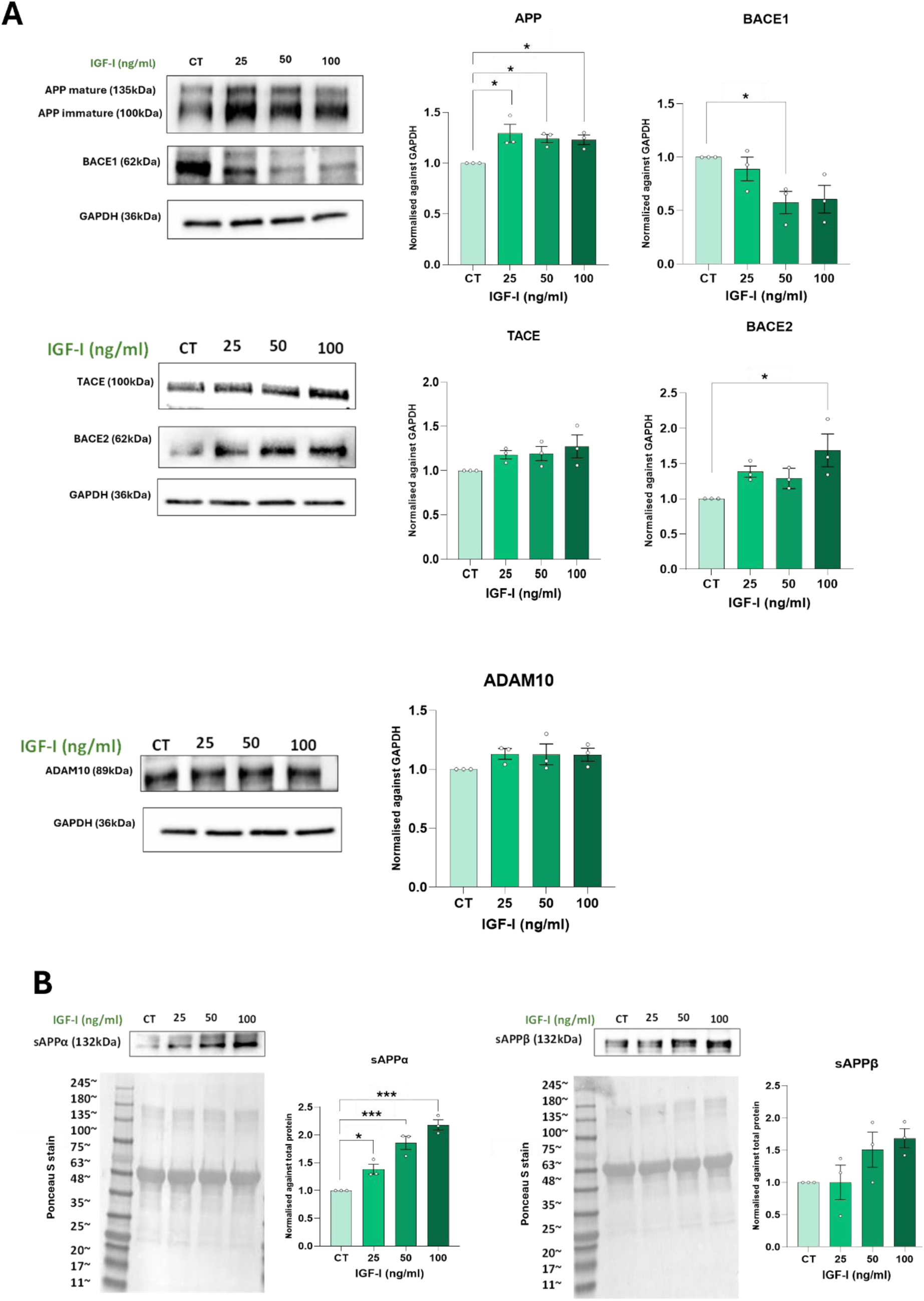
Assessment of the effects of IGF-I on APP and secretase enzymes in PC3 cells. **(A)** IGF-I caused a significant increase in abundance of both APP (p<0.05) and BACE2 (p<0.05). ADAM-10 and TACE were not significantly altered by IGF-I, whilst BACE1 was significantly reduced at the 50ng/ml IGF-I dose (p<0.05). **(C)** Assessment of sAPPα and sAPPβ secreted into the cell supernatants showed a dose-dependent increase in sAPPα which was significant at all three IGF-I doses (25ng/ml p<0.05, 50ng/ml p<0.001, 100ng/ml p<0.001). sAPPβ levels were also increased in response to IGF-I but not significantly so. Representative western blots are shown, and graphs represent densitometric analysis of bands corrected to either the GAPDH loading control or Ponceau S stain. Bars represent fold change from control (mean ± SEM, n=3).

#### Effects of the IGF1R inhibitor, AG1024, on APP and its cleavage in PC3 cells

In PC3 cells, AG1024, a selective IGF1R inhibitor, caused a reduction in APP abundance that was significant at the 3µM dose (p<0.05) and a reduction in the α-secretase BACE2 that was significant at the 2µM dose (p<0.05). ADAM-10 abundance was not significantly changed by AG1024 whilst TACE was significantly increased at the 3µM dose (p<0.05). There was a trend towards a dose-dependent increase in BACE1, but this did not reach statistical significance (**Fig 5A**). Assessment of secreted soluble APP fragments, sAPPα and sAPPβ, showed a reduction in sAPPα that was significant at the 2µM (p<0.01) and 3µM (p<0.01) doses, and a reduction in sAPPβ that was most significant at the lowest AG1024 dose (p<0.01) (**Fig 5B**).

**Figure 5:**
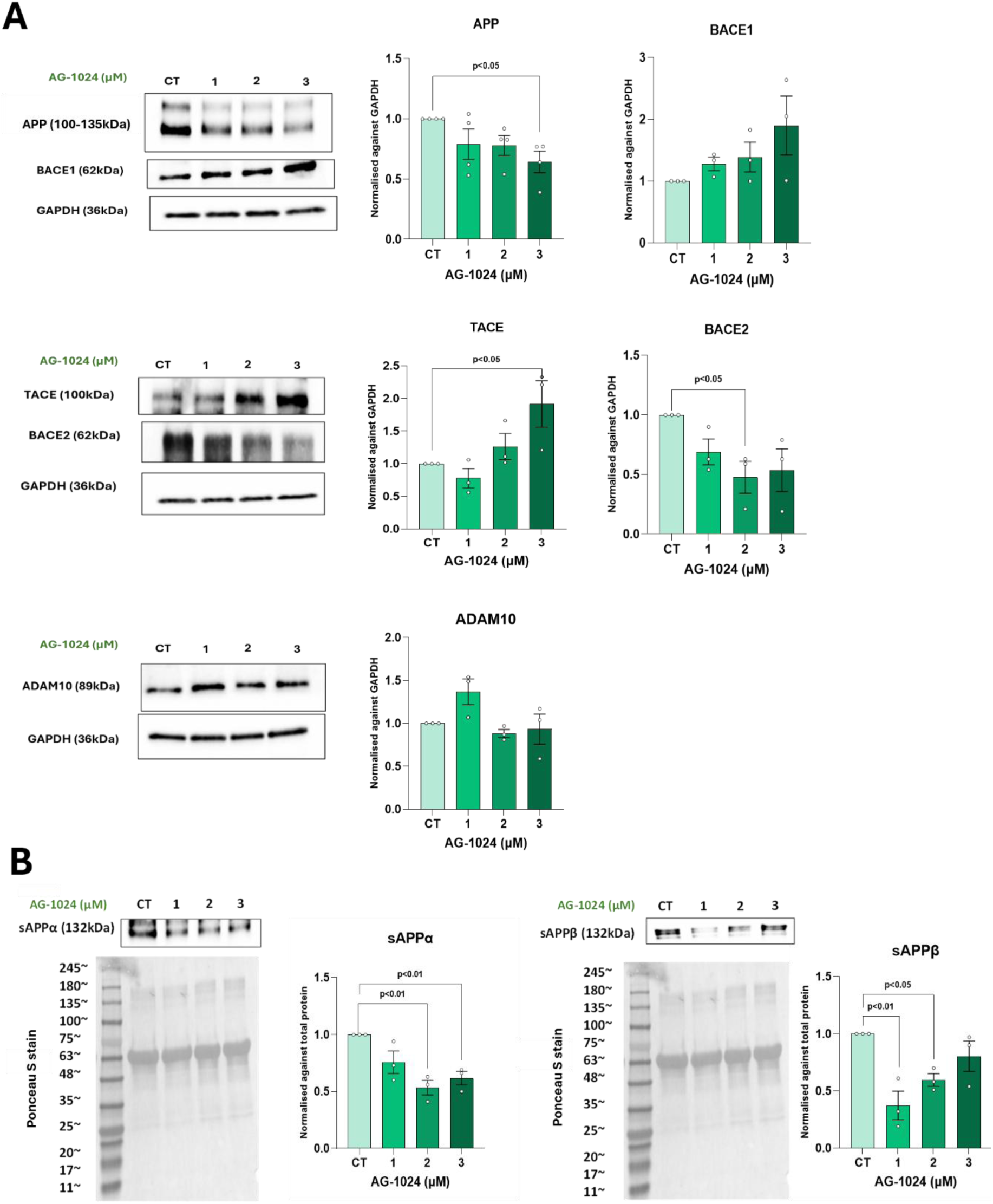
Assessment of the effects of AG1024 on APP and secretase enzymes in PC3 cells. **(A)** AG1024 caused a reduction in APP abundance that was significant at the 3µM dose (p<0.05) and a reduction in BACE2 that was significant at the 2µM dose (p<0.05). ADAM-10 abundance was not significantly changed by AG1024 whilst TACE was significantly increased at the 3µM dose (p<0.05). There was a trend towards a dose-dependent increase in BACE1, but this did not reach statistical significance. **(B)** Assessment of sAPPα and sAPPβ secreted into the cell supernatants showed a reduction in sAPPα that was significant at the 2µM (p<0.01) and 3µM (p<0.01) doses, and a reduction in sAPPβ that was most significant at the lowest AG1024 dose (p<0.01). Representative western blots are shown, and graphs represent densitometric analysis of bands corrected to either the GAPDH loading control or Ponceau S stain. Bars represent fold change from control (mean ± SEM, n=3).

#### Inhibiting α-secretases, ADAM-10 and TACE, effectively reduces sAPPα production, inhibits cell growth

We have shown that sAPPα production, produced by cleavage of APP by α-secretases ADAM-10 and TACE, is induced by key drivers of prostate cancer growth, androgen and IGF-I. A comparison of TCGA-PRAD and GTEx normal prostate datasets revealed a significant upregulation of *ADAM10* in tumour compared to normal tissues (**extended data Fig 1C**). Hence, we used a dual inhibitor of ADAM-10 and TACE, GW280264X (Tocris Bioscience), to assess the effects of inhibiting α-cleavage of APP in LNCaP and PC3 cells. Dose responses to GW280264X were first performed and showed a significant reduction in proliferation in both LNCaP and PC3 cells, but no significant effects in PNT2 cells (**Extended data Fig 2A-C**). Effective reduction of sAPPα production was confirmed by measurement of sAPPα in the cell supernatants. Further analysis of live and dead cell counts following GW280264X treatment in LNCaP and PC3 cells revealed a significant reduction in total cell counts in both LNCaP (**Extended data Fig 3A(i)**, p<0.05) and PC3 (**Extended data Fig 3B(i)**, p<0.05) cells when treated with 1000nM of inhibitor but had no effect on percentage viability in either (**Extended data Fig 3A(ii) and 3B(ii)**). A significant reduction in sAPPα secretion into the cell supernatant was confirmed at the 1000nM inhibitor dose in both cell lines (**Extended data Fig 3A(iii) and 3B(iii)**, p<0.05). Despite the growth inhibitory effects of sAPPα inhibition, it did not significantly enhance the effects of enzalutamide in LNCaP cells, AG1024 in PC3 cells or cabazitaxel in either cell line (**Extended data Fig 2C and D).**

#### sAPPα protects cells from the inhibitory effects of enzalutamide and AG1024 on cell viability

We have shown that key drivers of prostate cancer growth upregulate α-cleavage of APP and inhibiting α-secretase activity reduces cell proliferation. Next, we investigated the effects of addition of exogenous sAPPα to LNCaP and PC3 cells. Pre-incubation of LNCaP cells with 2nM sAPPα prior to enzalutamide (1µM) treatment, conferred enhanced survival (**Extended data Fig 3C**). Treatment with enzalutamide alone induced an 11% reduction in cell viability (p<0.001), whilst treatment with sAPPα and enzalutamide together showed only a 3% reduction in percentage cell viability (non-significant) compared to the untreated control. Cell viability was significantly increased in cells dosed with both sAPPα and enzalutamide compared to cells treated with enzalutamide alone (p<0.001). In PC3 cells, pre-incubation with 2nM sAPPα prior to AG1024 (2µM) treatment conferred survival, although cell death was still induced (**Extended data Fig 3D**). Cells treated with AG1024 alone showed a 16% reduction in cell viability (p<0.001), whilst treatment with sAPPα and AG1024 together showed a 10% reduction in percentage cell viability (p<0.001) compared to the untreated control. Cell viability was significantly increased in cells dosed with both sAPPα and AG1024 compared to cells treated with AG1024 alone (p<0.05).

#### Silencing APP induces apoptosis in prostate cancer but not normal cells

APP was successfully silenced in all cell lines **(Fig 6A, B s C(iii))**. APP silencing did not induce significant apoptosis in the normal PNT2 cells at either siRNA concentration **(Fig 6A(i-ii))**. In contrast, in LNCaP cells, there was a significant induction of apoptosis at both siRNA concentrations associated with a significant reduction in percentage live cells in the silenced population compared to the non-silencing control **(Fig 6B(i-ii))**, p<0.001)) and a significant increase in abundance of cleaved PARP (**Fig 6B(v-vi)**, p<0.01). In PC3 cells, the same pattern was observed: a significant reduction in percentage live cells compared to the non-silencing control (**Fig 6C(iii-iv)**) and a significant induction of apoptosis as indicated by increases in the abundance of cleaved PARP (**Fig 6C(v-vi)**, p<0.001).

**Figure 6:**
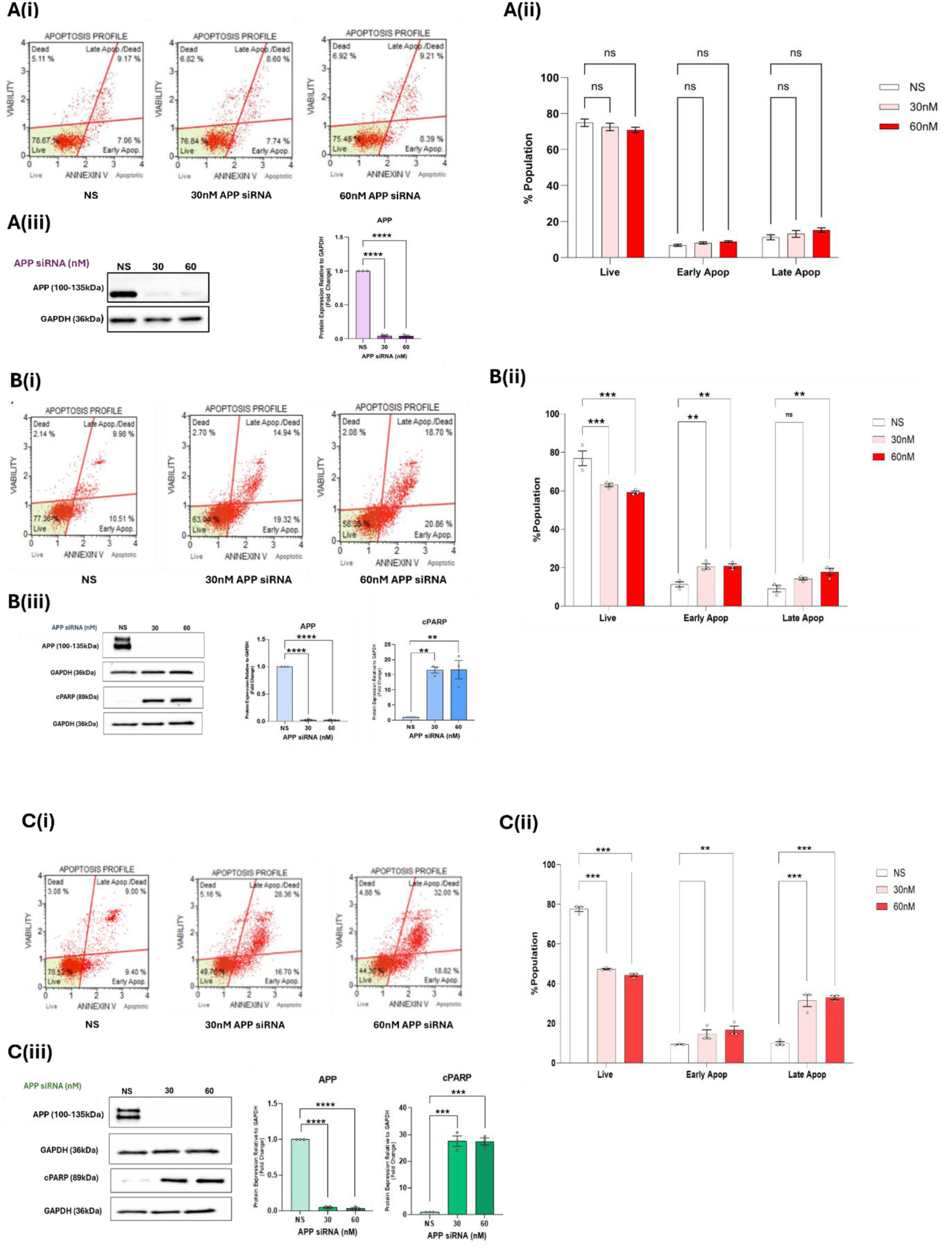
Effects of silencing the *APP* gene in (A) PNT2, (B) LNCaP and (C) PC3 cells. Results show representative cell apoptosis profiles **(i)**, percentage live and apoptotic cell populations **(ii)** and western blots with densitometric analysis to confirm APP silencing was successful and to assess apoptotic marker, cleaved PARP where appropriate **(iii)**. APP was successfully silenced in PNT2 cells at both siRNA concentrations (30 & 60nM) as shown by significant reduction in abundance of APP (**A(iii)**, p<0.001) but this did not induce a significant apoptotic response (**Ai-ii**). APP was successfully silenced in both LNCaP and PC3 cells at both concentrations of siRNA (30-60nM), as shown by significant reductions in the abundance of APP (LNCaP (**Biii)**, p<0.001; PC3 (**Ciii)**, p<0.001). *APP* silencing induced significant apoptosis in both LNCaP (B(i-ii); p<0.001) and PC3 (C(i-ii); p<0,001) cells. There was a significant increase in abundance of cleaved PARP in both LNCaP ((B(iii)); p<0.001) and PC3 ((C(iii)); p<0.001) cells. All graphs show mean ± SEM, n=3. For densitometric analysis, protein abundance was corrected to the GAPDH loading controls and bars represent fold change from the non-silencing control.

#### Effects of addition of IGF-I following APP silencing in PC3 cells

IGF-I is known to activate survival-promoting signalling pathways, including the PI3K/AKT pathway, contributing to resistance against chemotherapy, radiotherapy, and targeted therapies [25, 26]. We therefore anticipated that IGF-I would protect PC3 cells from APP silencing-induced apoptosis.

However, IGF-I treatment of PC3 cells did not protect cells from the apoptosis induced by silencing APP; apoptosis induction by APP silencing was similarly significant in both cells with or without IGF –I addition (**Fig 7A**; p<0.001) and cPARP levels were similar in both **(Fig 7B)**. There was also a concurrent loss of AKT signalling in *APP*-silenced cells, as shown by a significant reduction in AKT phosphorylation **(Fig 7B**, p<0.001), and addition of IGF-I did not restore signalling. To further understand the lack of IGF-I mediated survival and loss of AKT signalling, we assessed levels of the IGF1R and its signalling following *APP* silencing. Results showed a significant reduction in IGF1R abundance following APP silencing (**Fig 7C(i);** p<0.01). In LNCaP cells, *APP* silencing also resulted in a significant reduction in abundance of the AR (**Fig 7C(ii**); p<0.001), implying that androgen would also not induce survival effects in *APP* silenced cells.

**Figure 7:**
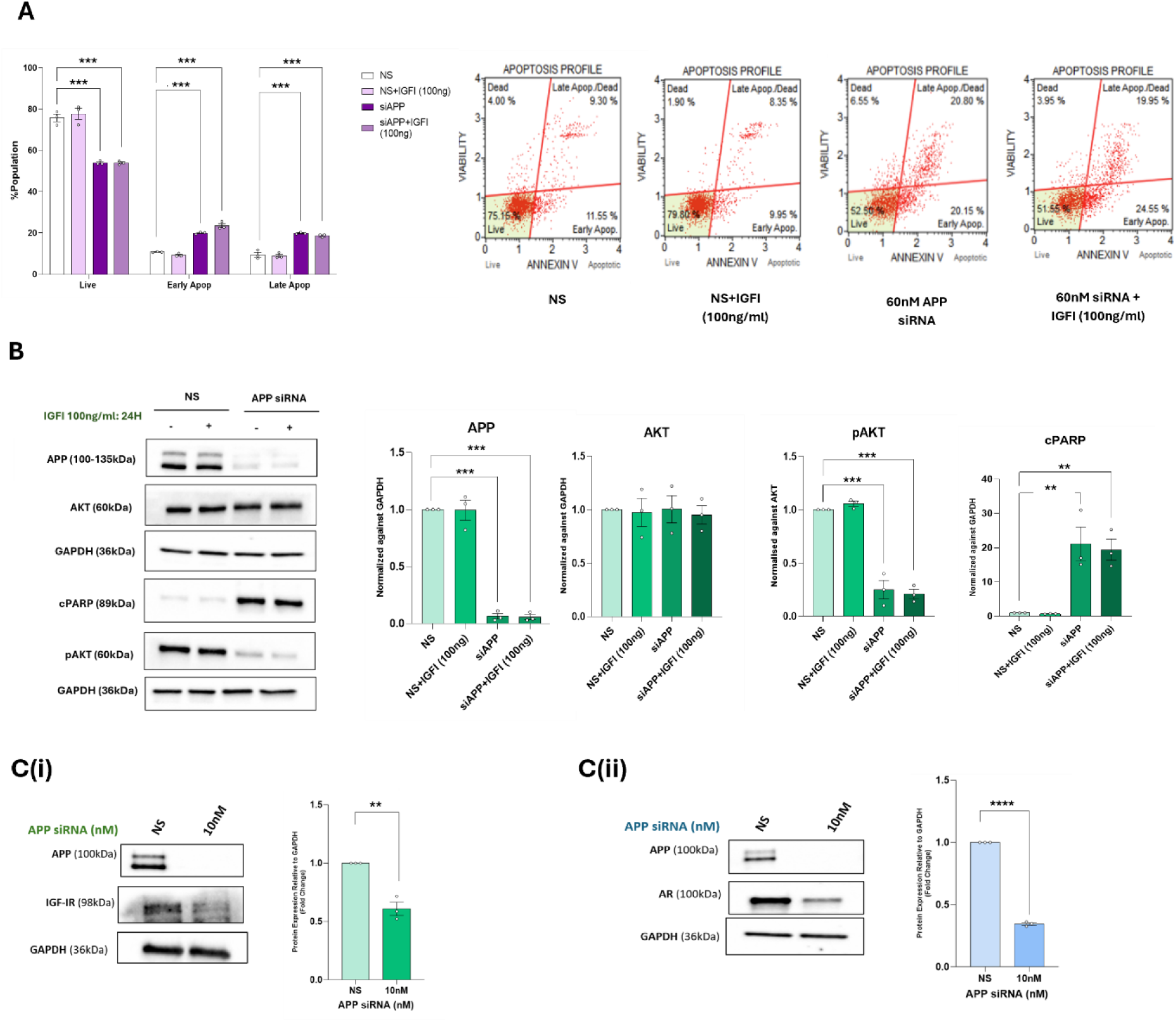
Effects of IGF-I addition following knockdown of APP. PC3 cells were transfected for 24 h with either non-silencing (NS) or APP siRNA (60nM) and then treated with 100ng/ml IGF-I for a further 24 h before assessment of apoptosis. **(A)** show apoptosis summary statistics and representative cell apoptosis profiles. IGF-I addition did not protect cells from induced apoptosis as significant apoptosis was apparent in silenced cells either with or without IGF-I addition (p<0.001). **(B)** shows western blots and densitometric analysis of apoptotic marker, cPARP, and downstream IGF-I signalling molecule AKT, where phosphorylation indicates IGF-I signalling. Silencing APP caused increased cPARP in cells either with or without IGF-I (p<0.001) and also resulted in loss of IGF-I signalling in both, as indicated by significantly reduced phosphorylated AKT (pAKT; p<0.001). **(C)** *APP* silencing caused a significant reduction the IGF1R in PC3 cells (**(i);** p<0.01) and in AR in LNCaP cells ((**ii**); p<0.001). Representative western blots are shown. Graph*s* represent densitometric analysis, mean ±SEM, n=3.

#### APP positively correlates with the AR and the IGF1R in prostate tumour tissue

To further understand the relationship between APP and AR and the IGF1R, we assessed correlations in immunohistochemical staining in tumour tissue from the PrEvENT study (**Fig 8A)**. Results showed positive correlations between APP and both AR and the IGF1R; unlike the statistically significant positive correlation between APP and the IGF1R (**Fig 8B(i)** r=0.36, p=0.001), the positive correlation between APP and AR did not reach statistical significance in this small cohort (**Fig 8B(ii)**; r=0.19, p=0.08) We also assessed correlations between genes expressing APP, AR and IGF1R in the TCGA-PRAD dataset. Again, significant positive correlations were found between *APP* and the expression of both the IGF1R (**Fig 8C(i)**; r=0.33, p<0.0001) and the AR (**Fig 8C(ii)**; r=0.27, p<0.0001).

**Figure 8:**
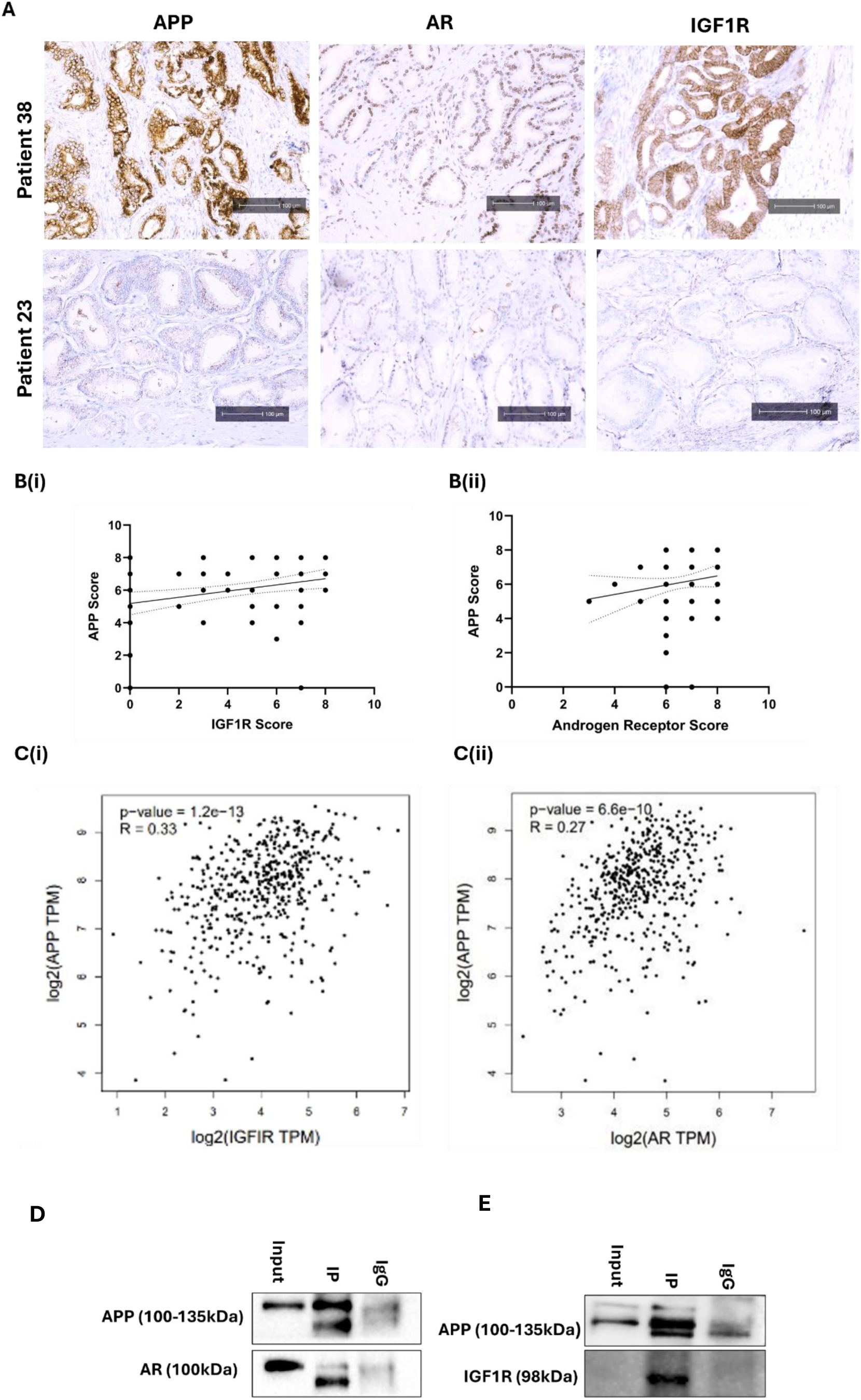
Correlations between APP and AR and the IGF1R in prostate tumour tissue and APP co-immunoprecipitation. **(A)** Representative immunohistochemical staining of APP, AR and the IGF1R in prostate tumour tissue from two participants in the PrEvENT study, one with high abundance and one with low abundance of all three proteins. **(B)** Correlation between APP and AR **(i)** and APP and the IGF1R **(ii**) in the immunohistochemical staining of the PrEvENT study (n=96). Results showed a non-significant positive correlation between APP and AR (r=0.19, p=0.08), whilst APP and the IGF1R were significantly positively correlated (r=0.36, p=0.001). **(C)** Correlation between *APP* and *AR* expression (**i**) and *APP* and *IGF1R* expression (**ii**) in the TCGA-PRAD cohort (n=452). There was a significant positive correlation between *APP* and *AR* (r=0.27, p<0.0001) and between *APP* and *IGF1R* (r=0.33, p<0.001). **APP co –immunoprecipitation.** Co-immunoprecipitation in **(D**) LNCaP and **(E)** PC3, confirmed by APP bands in the input and co-IP lanes, but reduced or no signal in the IgG isotype control lane. Blots also show the presence of AR in the co-IP lane in LNCaP cells and the IGF1R in the co-lane in PC3 cells, confirming APP interaction with these receptors. Representative western blots are shown. N=3.

#### APP interacts with AR and IGF1R

To confirm if an interaction existed between APP and the AR in LNCaP cells and between APP and the IGF1R in PC3 cells, we performed co-immunoprecipitation. Western blots confirmed that, in LNCaP cells, AR was present within the pulled-down APP complex and the IGF1R was present in the complex in PC3 cells (**Fig 8 D and E** respectively).

#### Pathways associated with APP silencing

To further understand the effects of *APP* silencing, we performed proteomic analysis. In LNCaP cells, *APP* silencing resulted in 4 significantly upregulated proteins and 28 significantly downregulated. **Extended data table 1** summarises the upregulated and 10 most downregulated proteins and their functions. In PC3 cells, *APP* silencing caused a significant increase in the abundance of 4 proteins, while 19 were decreased. **Extended data table 2** summarises the upregulated and 10 most downregulated proteins and their functions. Volcano plots highlighting the significantly up –and down-regulated proteins in LNCaP and PC3 cells are shown in **Extended data figure 4A and B**.

Gene ontology (GO) analysis was performed to understand the pathways affected by *APP* silencing. In LNCaP cells, GO enrichment revealed that the downregulated proteins were associated with RNA splicing and ribosomal biogenesis, whereas the upregulated proteins were linked to apoptotic signalling pathways (**Extended data figure 4C and D).** In PC3 cells, GO enrichment showed that the downregulated proteins were primarily involved in carbohydrate metabolism and cell cycle regulation, whereas the significantly upregulated proteins were again associated with apoptosis and also with wound healing (**Extended data figure 4E and F**).

## Discussion

This study provides seminal evidence to show that APP may be an Achilles’ heel to exploit in the treatment of prostate cancer. Performing tissue studies, we confirmed that APP is a clinically relevant target in prostate cancer. In a tissue cohort from 96 men with localised prostate cancer, we confirmed a significant increase in the abundance of APP in tumour compared to surrounding benign tissue, that has been described previously [11]. Within the tumour samples we found a trend towards reduced APP with advancing Gleason grade, a correlation that was significant when analysing *APP* gene expression levels in the TCGA PRAD dataset. This trend would be consistent with APP being an androgen-regulated gene [11] and of particular importance in lower grade, androgen sensitive prostate cancer. Given that our tissue cohort held limited numbers of higher Gleason grade tissue it would be of interest to investigate this further in high grade and metastatic samples to confirm these findings.

Our study investigated the regulation of APP by key drivers of prostate cancer development, androgen and IGF-I. With LNCaP cells, androgen increased APP levels and promoted α-secretase processing of APP, giving rise to increased sAPPα production. Previous studies have shown androgen regulation of APP in both prostate and breast cancer cells [11, 14] and, in prostate cancer cells, upregulation of α-secretase, *ADAM10* expression [27, 28], as well as an increase in ADAM10 protein levels and its nuclear translocation in response to androgen [29]. Whilst one of these studies showed that a combination of androgen and IGF-I treatment upregulated ADAM10 in LNCaP cells [27], to our knowledge no previous studies have examined the effects of IGF-I on APP and its processing in prostate cancer cells.

Because IGF-I has been associated with more aggressive prostate cancer [17], we explored its effects on APP in the androgen independent cell line, PC3. IGF-I caused an increase in APP levels, and despite increases in the α-secretases being largely non-significant, an increase in sAPPα was still seen. Interestingly, sAPPβ levels were also increased in response to IGF-I, despite BACE1 being decreased.

It is well established that androgen and IGF signalling pathways interact, with AR signalling enhancing IGF signalling by increasing transcription of the IGF1R and enhancing IGF-I secretion through inhibition of IGF binding protein-3 [30]. Future investigation, therefore, of combined androgen and IGF-I effects on APP in androgen sensitive prostate cancer may be of benefit and offer novel ways to target both androgen and IGF signalling. Given that both APP and sAPPα have been associated with cell growth [15, 31], our data provide a mechanism by which DHT and IGF –I promote cell growth and survival of prostate cancer cells.

Notably, correlation analysis of the TCGA PRAD dataset showed significant positive associations between APP and both the AR and the IGF1R, and these associations were confirmed at the protein level in our smaller immunohistochemical study. Our co-immunoprecipitation experiments further indicated that an interaction existed between these proteins, which is particularly evident with APP and the IGF1R. This may offer future opportunities for more targeted approaches to IGF1R inhibitor therapies, which have so far proved challenging, largely due to a lack of predictive biomarkers [25].

Importantly, when we silenced *APP* in LNCaP and PC3 cells, there was a significant induction of apoptosis which was not seen in normal prostate PNT2 cells, again implying that APP is vital for growth and survival of prostate cancer cells. Perhaps predictably, given the prior evidence for interactions between APP and both the AR and the IGF1R, we also showed that levels of these receptors were significantly reduced following *APP* silencing and, hence IGF-I was unable to protect cells from *APP* silencing-induced apoptosis. Proteomic analysis following *APP* silencing revealed upregulation of proteins involved in apoptosis pathways and downregulation of proteins involved in cell cycle regulation and RNA processing. Additionally, some proteins that function in cell survival and cancer growth were also upregulated, consistent with induction of stress –response mechanisms. Interestingly, there were several proteins that were significantly downregulated (RHPN2, DICER1, FYCO1, C19orf53, PLEKHA1, CHCHD10, TNRC18, AAR2 and CDC25) and one protein (TNFRSF12A) that was significantly upregulated in both cell lines following *APP* silencing. Whilst these proteins may indicate activated pathways and processes that are common across the cell lines following *APP* silencing, examination of proteins that are altered differently in the two cell lines may offer further insight into APP’s role in different contexts and different disease stages. For example, ribosomal biogenesis factor (RBIS) was significantly downregulated in LNCaP but not PC3 cells. Ribosome biogenesis is essential for cell growth and proliferation and has been associated with cancer metastasis and therapy resistance [32]. Whilst ribosome biogenesis may be a target for treatment of CRPC [33], it is also regulated by androgen, therefore may also be of importance in androgen-sensitive prostate cancer [34]. RBIS has been shown to correlate with tumour stage, Gleason grade and poor progression-free survival [35], hence its downregulation following *APP* silencing may offer favourable outcomes for prostate cancer patients. In PC3 cells, *APP* silencing caused significant downregulation of connexin43 (encoded by the *GJA1* gene) which was not seen in LNCaP cells. Connexin43 is a gap junction protein that is often downregulated in the early stages of prostate cancer, however specific isoforms such as GJA1-20k have been shown to be upregulated in advanced bone-metastasizing prostate cancer [36, 37]. These results provide important initial insights into APP’s mechanism of action in prostate cancer and identify additional molecules of interest that require further investigation to fully understand its interactions.

It is also vital to understand the effects of the cleavage products of APP in cancer, given that these have been shown to have differing functions. For example, non-amyloidogenic cleavage of APP by α-secretases produces sAPPα which has been shown to exhibit neuroprotective effects in AD models [38], as well as promoting growth in breast and prostate cancer models [11, 15]. Conversely Aβ, produced by BACE1 cleavage of APP, is associated with neuronal death in AD and has also been shown to inhibit growth of cancer cells [39]. Hence, differential regulation of APP processing in the two diseases may offer one mechanism underlying the inverse association between the two: upregulated β-cleavage in AD promoting cell death and upregulated α-cleavage in cancer promoting cell growth and survival. APP cleavage may also be influenced by its isoform status, the three most abundant isoforms being APP_695_, APP_751_ and APP_770_ [5, 40]. Whilst APP_695_ is the most abundantly expressed isoform in neuronal cells, the other two isoforms are more abundant in non-neuronal and peripheral cells [5]. Previous studies in models of AD have indicated differential isoform expression both in different cell types and in different subcellular locations, which suggests differing functions and effects [40, 41]. Investigation of APP isoforms and their differential effects in cancer cells have not previously been conducted but would be of great benefit in further understanding the role of APP in cancer and may identify a novel, more specific therapeutic target.

To further examine the effects of sAPPα on prostate cancer cells, we treated cells with exogenous sAPPα and inhibited its endogenous production using a dual ADAM-10/TACE inhibitor. Whilst addition of sAPPα alone did not significantly affect cell proliferation in our models, it did have an impact on survival and, notably, reduced the efficacy of enzalutamide in LNCaP cells and AG1024 in PC3 cells. Consistent with a survival effect of sAPPα, inhibiting endogenous sAPPα production also led to significant reductions in cell proliferation in both LNCaP and PC3 cells but did not significantly affect normal PNT2 cells. The effect in cancer cells appeared to be growth reducing rather than an induction of cell death which was seen with *APP* silencing, indicating that full-length APP and sAPPα affect cancer cells by different mechanisms. Whilst sAPPα inhibition caused a significant reduction in proliferation, it did not significantly enhance the effects of enzalutamide, IGF1R inhibition or chemotherapy.

Given that the effects of *APP* silencing and α-secretase inhibition have significant impact in cancer and not normal prostate cell lines, both offer exciting new therapeutic targets for prostate cancer treatment. Drugs that reduce APP translation have been previously developed for AD treatment [42–44] and indeed some have entered clinical trials for treatment of AD and Parkinson’s disease [45, 46]. These may represent new opportunities for drug repurposing for prostate cancer, either as standalone treatments or in combination with current therapies. Targetting of α-secretase enzymes, including ADAM-10 and TACE for cancer therapy has already been considered [47–49], but, to our knowledge, not in the context of APP cleavage. ADAMs cleave a broad range of ligands and receptors, including molecules of particular importance in cancer, for example, Notch, epithelial growth factor receptor (EGFR), cytokines including tissue necrosis factor (TNF)-α and adhesion molecules [50]. Previous ADAM targetting strategies have included low molecular weight synthetic inhibitors [51], purified ADAM-10 pro-domain [52] and monoclonal antibodies [53]. One low molecular weight inhibitor, INCB3619, showed promise in *in vitro* models of non-small cell lung carcinoma (NSCLC) [54] and breast cancer [55]. A second-generation inhibitor, INCB7839, has shown promising results in a Phase I/II trial in HER2+ breast cancer, improving response to trastuzumab in a subpopulation of HER2+ patients [56]. Challenges of ADAM targetting for cancer therapy, however, include redundancy among ADAM family members and non-tumour effects and toxicity, given their important functions in normal tissues [50]. Given these challenges, targetting APP directly may offer a more effective and safer alternative to ADAM inhibition.

## Conclusions

Notably, we have shown that APP is overexpressed in prostate tumour tissue and its processing is regulated by key drivers of prostate cancer growth, androgen and IGF-I, promoting non-amyloidogenic cleavage of APP. The resulting sAPPα peptide is associated with cell survival and reduced sensitivity to enzalutamide, whilst inhibition of sAPPα production *via* an ADAM-10/TACE dual inhibitor causes reduced cell growth. Silencing the *APP* gene causes significant apoptosis in prostate cancer but not normal prostate cells, indicating that APP might offer an attractive therapeutic target for prostate cancer treatment. Even more exciting is that drugs that target APP, for example buntanetap, are already in clinical trials for those with early AD (e.g NCT06709014), hence may offer an exciting and feasible approach for drug repurposing in the treatment of prostate cancer.

## Ethics approval and consent to participate

n/a

## Consent for publication

n/a

## Data availability

The proteomic data generated and analysed during the current study are available from the corresponding author on reasonable request.

## Authors’ contributions

RB, EG and MA data acquisition and analysis. RB and CP drafted the manuscript. CP conceived and supervised the work. All authors read and contributed to the final version of the manuscript.

## List of Abbreviations

AD: Alzheimer’s Disease
ADAM: A Disintegrin and Metalloproteinase
AKT: Protein Kinase B
APP: Amyloid Precursor Protein
AR: Androgen Receptor
Aβ: Amyloid-beta
BACE: Beta-site APP Cleaving Enzyme
BrdU: 5-Bromo-2’-Deoxyuridine
cPARP: Cleaved Poly(ADP-ribose) Polymerase
CRPC: Castrate-Resistant Prostate Cancer
DHT: Dihydrotestosterone
EGFR: Epidermal Growth Factor Receptor
FFPE: Formalin-Fixed Paraffin-Embedded
IGF: Insulin-like Growth Factor
MAPK: Mitogen-Activated Protein Kinase
PARP: Poly(ADP-ribose) Polymerase
PI3K: Phosphoinositide 3-Kinase
PrEvENT: Prostate Cancer: Evidence of Exercise and Nutrition Trial
sAPP: Soluble Amyloid Precursor Protein
TACE: TNF-α Converting Enzyme
TCGA: The Cancer Genome Atlas
TNF-α: Tumour Necrosis Factor Alpha
TTI: Tritiated Thymidine Incorporation (Assay)

## Acknowledgements

We would like to thank participants of the Prostate Cancer: Evidence of Exercise and Nutrition Trial (PrEvENT) and those involved in conceptualisation of the trial, in particular Prof Richard Martin, Prof Athene Lane and Dr. Lucy Hackshaw-McGeagh. We would also like to acknowledge Southmead Charity Trust for supporting this work.

