## Supplementary data for "Amyloid precursor protein: an Achilles’ heel to exploit in the treatment of prostate cancer"

Extended data figures

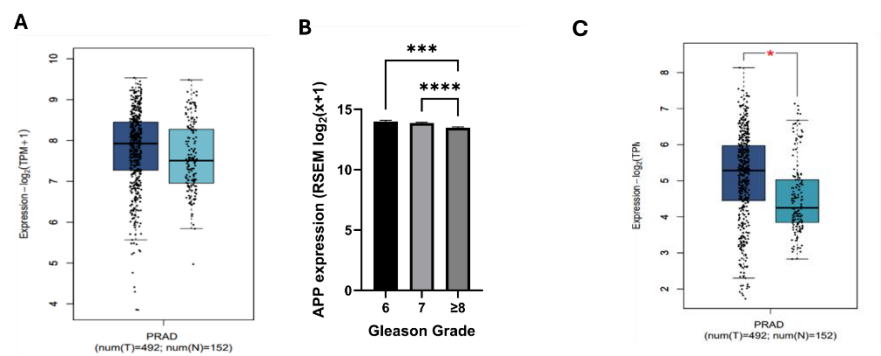

**Extended data figure 1: Assessment of APP and ADAM10 gene expression in human prostate tissue.** mRNA sequencing data from TCGA (tumour; n= 497) and GTEx (normal; n=152) datasets shows (A) a non-significant increase in APP expression in tumour tissues compared to normal. (B) In the TCGA cohort there was a significant reduction in APP expression in Gleason grades of 8 higher compared to both Gleason grade 6 (p<0.001) and Gleason grade 7 (p<0.0001) (Gleason 6 n=45, Gleason 7 n=247, Gleason 8+ n=205). (C) mRNA sequencing data from TCGA datasets also shows a significant (P<0.05) increase in ADAM-10 expression in tumour tissues compared to normal.

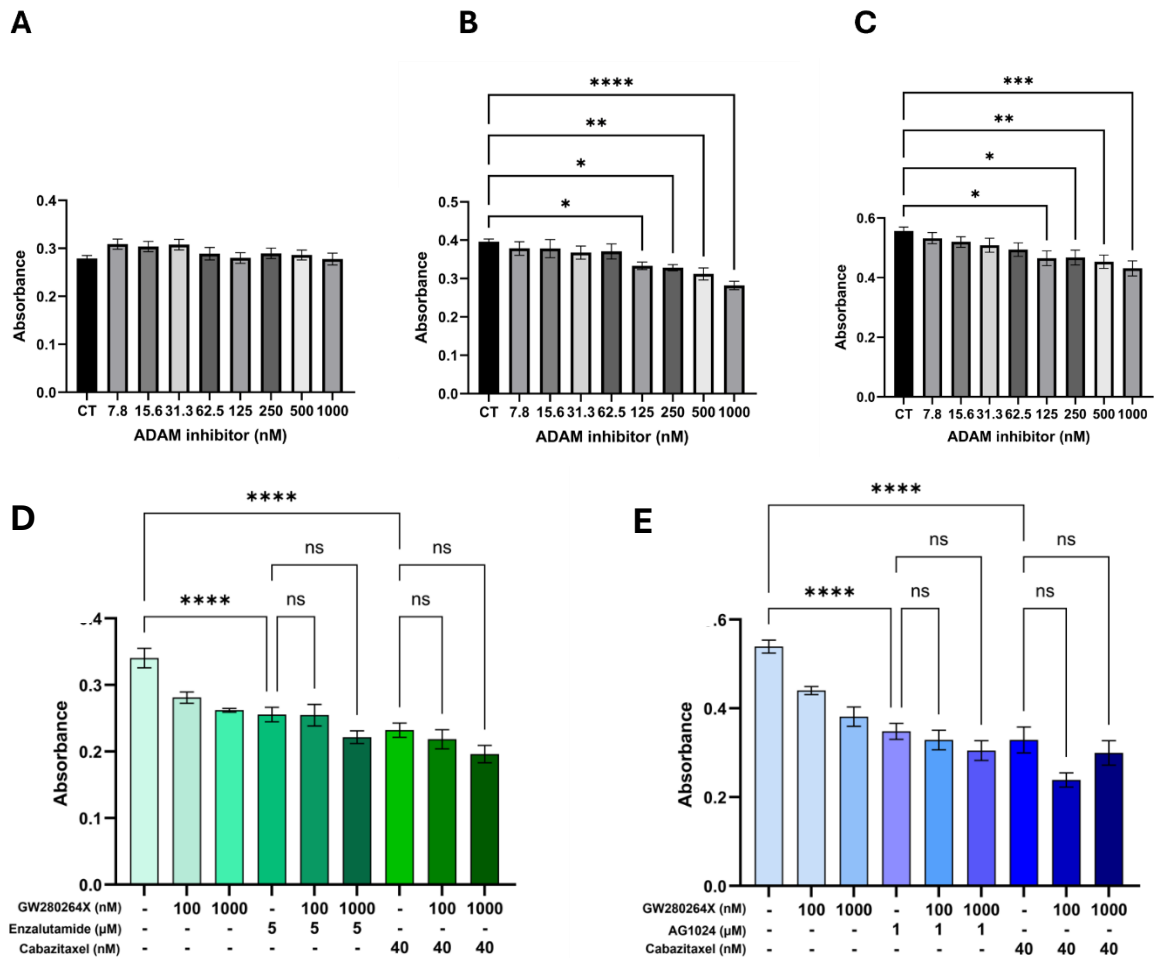

**Extended data figure 2: Dose response to ADAM inhibitor, GW 280264X and its effect on the sensitivity to enzalutamide, AG1024 and cabazitaxel.** PNT2 (A), LNCaP (B) and PC3 (C) cells were treated with ADAM inhibitor (0-1000nM) for 48 h and proliferation assessed by BrdU incorporation assay. ADAM inhibition did not significantly affect proliferation in PNT2 cells but caused significant, dose-dependent reduction proliferation in both LNCaP and PC3 cells. (D) In LNCaP cells, ADAM inhibitor GW280264X did not significantly affect the efficacy of enzalutamide or cabazitaxel. (E) In PC3 cells ADAM inhibitor GW280264X did not significantly affect the efficacy of IGF1R inhibitor AG1024 or cabazitaxel. Graphs represent mean ±SEM, n=3.

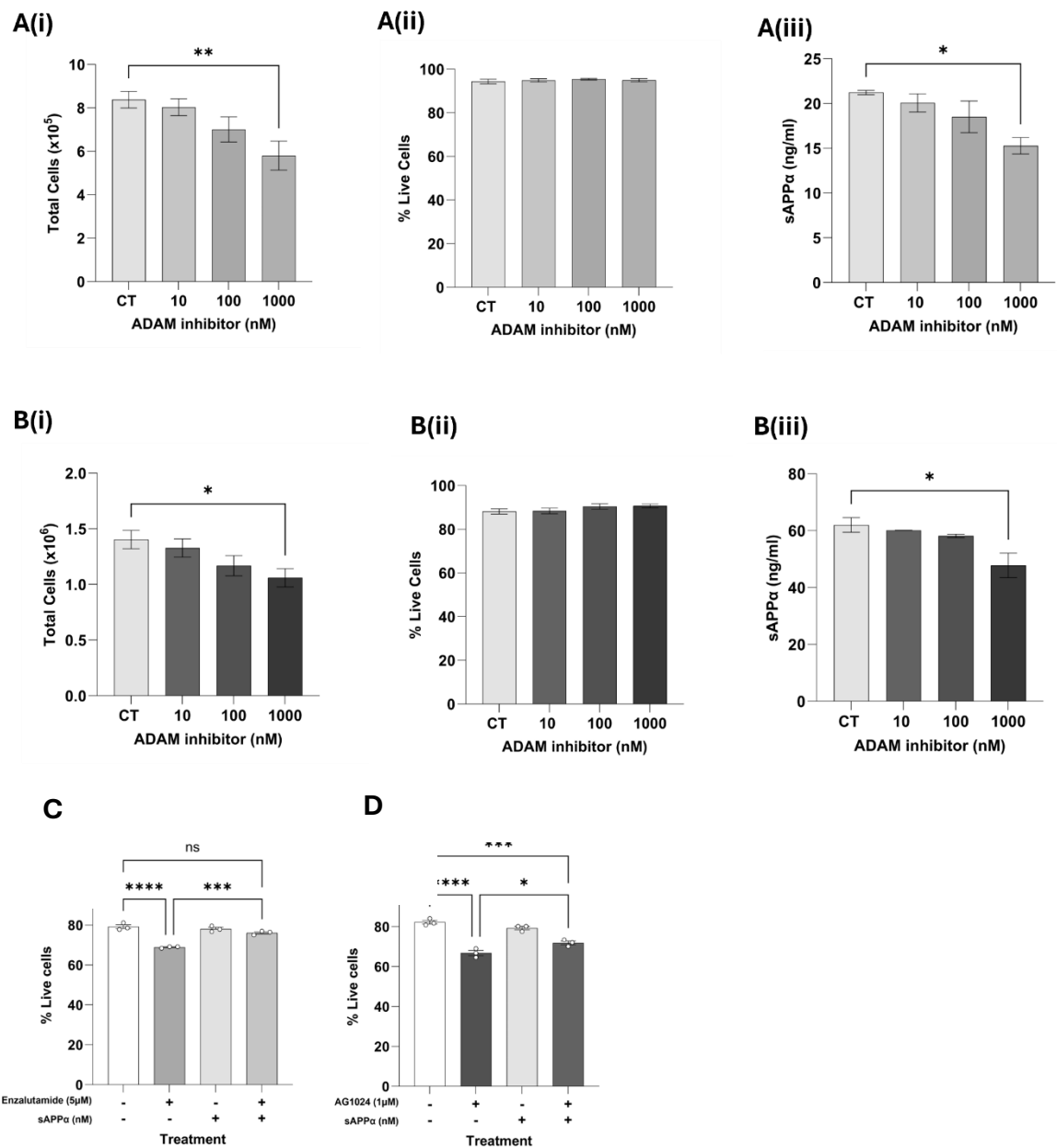

**Extended data figure 3: Effect of α-secretase inhibition in LNCaP and PC3 cells and the effect of pre-treatment with sAPPα on the response to enzalutamide in LNCaP and AG1024 in PC3 cells.** LNCaP (A) and PC3 (B) cells were incubated with GW280264X, a dual inhibitor of ADAM-10 and TACE, for 48 h then live and dead cells were counted and levels of secreted sAPPα were measured. (i) shows total cell counts, (ii) shows percentage cell viability and (iii) show levels of sAPPα in the cell supernatants. ADAM inhibition caused a significant reduction in total cells at the 1000nM dose ($p<0.05$) but did not affect percentage viability. Secretion of sAPPα was also significantly reduced at the 1000nM dose in both cell lines ($p<0.05$). Bars represent mean ±SEM. N=3. (C) Pre-incubation of LNCaP cells with 2nM sAPPα before treatment with 1μM enzalutamide significantly increased percentage viability compared to cells treated with enzalutamide alone ($p<0.001$). (D) Pre-incubation of PC3 cells with 2nM sAPPα before treatment with 2μM AG1024 significantly increased percentage viability compared to cells treated with enzalutamide alone ($p<0.05$). Bars represent mean ±SEM. N=3.

| Level | Protein Name | Function |
| --- | --- | --- |
| **Upregulated** | **SEC11C** | Protein processing in the ER |
|  | **TNFRSF12A** | TNF receptor; promotes survival and migration |
|  | **GDF15** | Stress cytokine; regulates tumour growth |
|  | **PARD6B** | Control cell polarity and junction assembly |
| **Downregulated** | **RBIS** | Ribosomal protein; protein synthesis |
|  | **RHPN2** | Cytoskeleton regulation and migration |
|  | **FYCO1** | Autophagosome transport |
|  | **DICER1** | miRNA processing |
|  | **PPFIA2** | Chromatin remodeling |
|  | **C19orf53** | Cell cycle regulation |
|  | **ZNF593** | Transcription regulation |
|  | **DDX21** | RNA processing and ribosomal biogenesis |
|  | **TMB1M6** | Intracellular trafficking |
|  | **CDC25C** | Cell cycle control, drives mitosis |

Extended data Table 1: Most significantly up- and downregulated proteins following *APP* silencing in LNCaP cells.

| Level | Protein Name | Function |
| --- | --- | --- |
| **Upregulated** | **LDLR** | cholesterol uptake; promotes survival and proliferation |
|  | **TNFRSF12A** | TNF receptor; promotes survival and migration |
|  | **TMEM209** | Protein transport |
|  | **S100P** | Calcium-binding protein; cancer proliferation and invasion |
| **Downregulated** | **CHCHD10** | Mitochondria protein |
|  | **RHPN2** | Cytoskeleton regulation and migration |
|  | **FYCO1** | Autophagosome transport |
|  | **DICER1** | miRNA processing |
|  | **PLEKHA1** | Binds to PIP3; promotes signaling pathway PI3K/AKT |
|  | **C19orf53** | Cell cycle regulation |
|  | **PPP4R3B** | Regulates DNA damage repair and cell cycle progression |
|  | **STYX** | Regulate signaling pathways MAPK/ERK |
|  | **KDM1B** | Epigenetic regulator; controls gene expression |
|  | **CDC25C** | Cell cycle control drives mitosis |

Extended data Table 2: Most significantly up- and downregulated proteins following *APP* silencing in LNCaP cells.

A

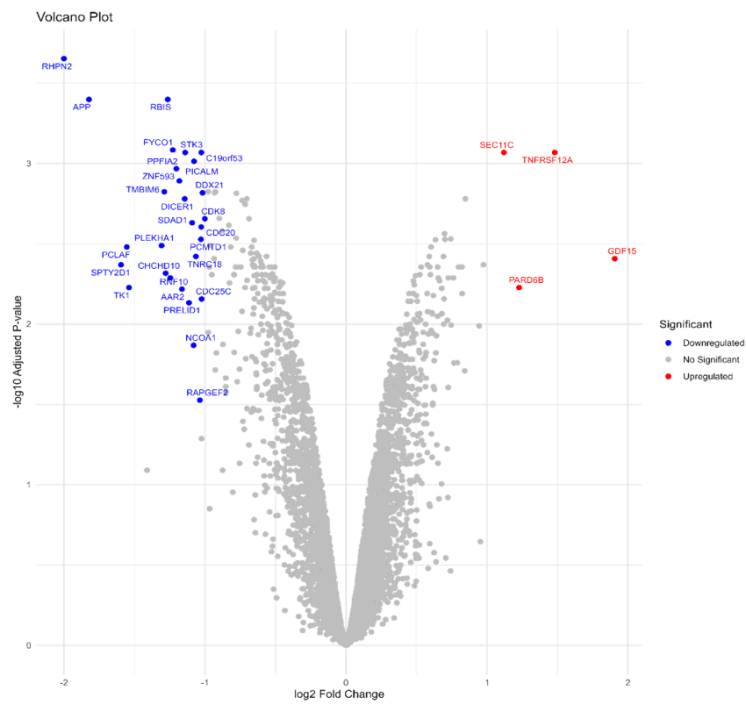

B

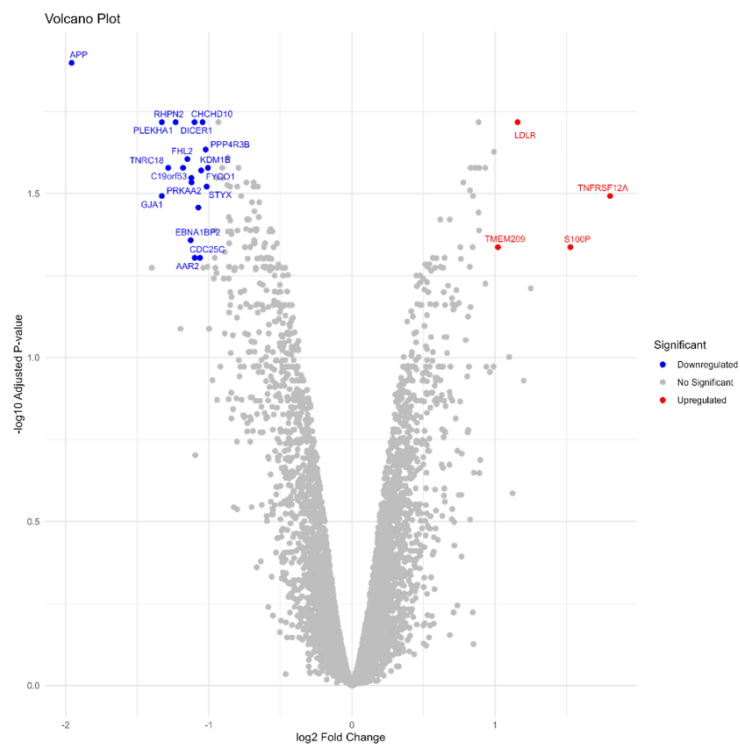

50

51

52

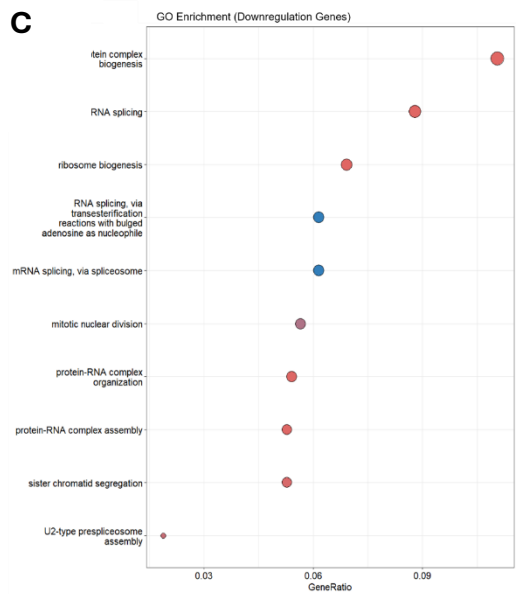

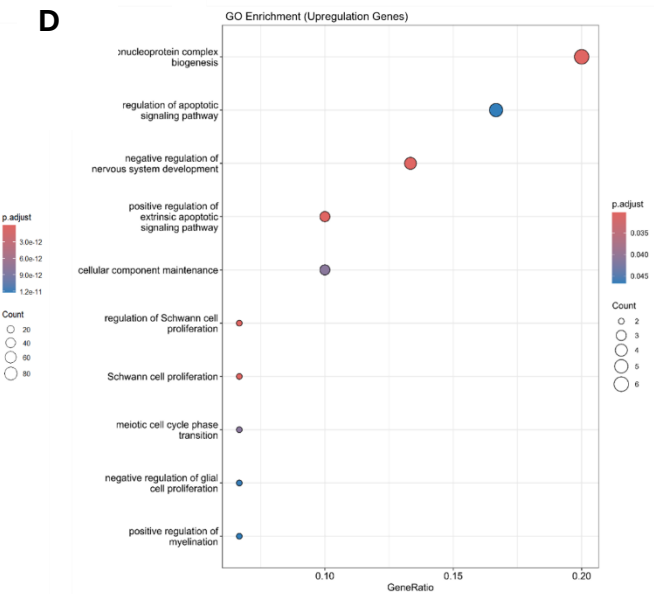

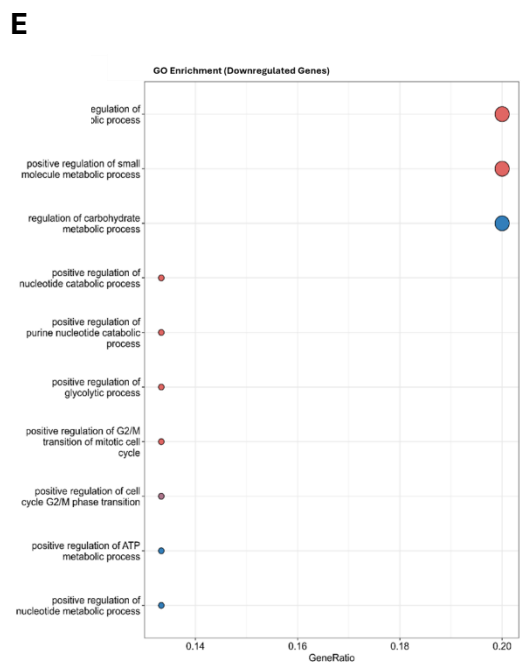

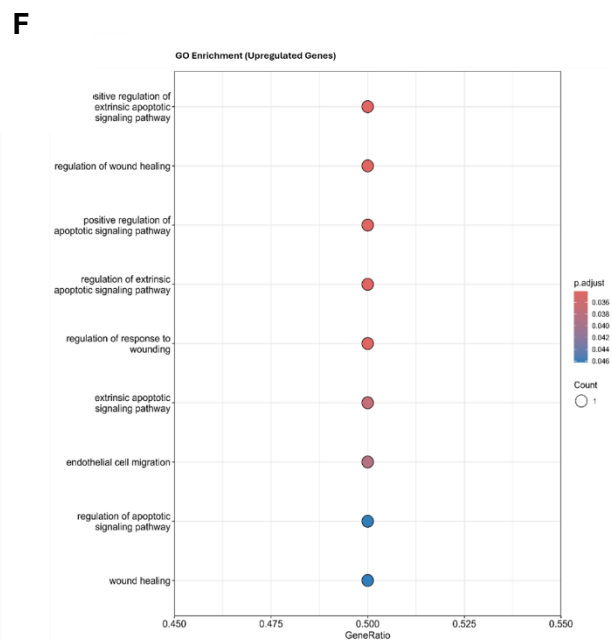

**Extended data figure 4.: Significantly up- and downregulated proteins following *APP* silencing.** TMT proteomic analysis was performed following *APP* silencing. Volcano plots show most upregulated (red) and downregulated (blue) proteins in LNCaP (A) and PC3 (B) cells. N=3. **GO enrichment analysis of proteomic data following *APP* silencing.** (C) shows downregulated and (D) shows upregulated processes in LNCaP cells. Most downregulated pathways included RNA splicing and ribosome biogenesis. Significantly upregulated processes included apoptotic signalling pathways. (E) shows downregulated and (F) shows upregulated processes in PC3 cells. The most significantly downregulated pathways include metabolic regulation. The most significantly upregulated processes include apoptotic signalling a wound healing regulation.
